# Predicting the Influence of Fat Food Intake on the Absorption and Systemic Exposure of Small Drugs using ANDROMEDA by Prosilico Software

**DOI:** 10.1101/2022.12.05.519072

**Authors:** Urban Fagerholm, Sven Hellberg, Jonathan Alvarsson, Ola Spjuth

## Abstract

**Introduction:** The ANDROMEDA software by Prosilico has previously been successfully applied and validated for predictions of absorption characteristics of small drugs in man. The influence of fat food on the gastrointestinal uptake and systemic exposure of drugs have, however, not yet been evaluated with the software.

**Objective and Methodology:** The main objective was to use ANDROMEDA to predict area under the plasma concentration-time curve ratios in the fed (fat food) and fasted states (AUC_fed_/AUC_fast_) for small drugs (including those marketed in 2021) and compare results with corresponding measured clinical estimates. Actual dose sizes were considered. Another objective was to compare the performance of ANDROMEDA *vs* physiologically based pharmacokinetic (PBPK) modelling and simulations by The Food Effect PBPK IQ Working Group. PBPK results generated using Simcyp and GastroPlus software were based on various physicochemical, *in vitro* and *in vivo* data and a decision tree for model verification and optimization.

**Results and Discussion:** 63 drugs, including 17 new drugs, with observed AUC_fed_/AUC_fast_ between 0.2 and 5.5 were found and used for this evaluation. Predicted AUC_fed_/AUC_fast_ had mean and maximum errors of 1.5- and 4.1-fold, respectively, and the predictive accuracy (correlation between predicted and observed AUC_fed_/AUC_fast_; Q^2^) was 0.3. 14 % of predictions had >2-fold error. For 72 % of drugs, food interaction class was correctly predicted. The level of predictive accuracy was overall similar to results obtained with PBPK modelling and simulations, however, with lower maximum error and higher compound coverage. With PBPK models, maximum simulation error was 7.7-fold and 3 highly lipophilic compounds were not possible to simulate.

**Conclusion:** The results validate ANDROMEDA for prediction of fat food-drug interaction size for small drugs in man. Major advantages with the methodology include that prediction results are produced directly from molecular structure and oral dose and are similar to PBPK-simulation results obtained using *in vitro* and clinical data. Furthermore, ANDROMEDA showed lower maximum errors and wider compound range.

## Introduction

When taken together with drugs, fat food may change their gastrointestinal uptake via changes in motiliy and transit time (reduced time for absorption of compounds with limited permeability (P_e_)) and/or solubility/dissolution (generally enhanced for lipophilic compounds with limited solubility/dissolution) (Deng et al. 2017). Mechanisms of food effects on gastrointestinal drug solubility/dissolution include increased bile acid and phospholipid concentrations, changes in pH and viscosity, and ion-pairing. Food might also change the oral bioavailability of drugs via changes in splanchnic and hepatic blood flows (such as for metoprolol) and lymphatic uptake (such as for venetoclax) (Emami Riedmaier et al. 2020).

It has been estimated that conmittant intake of food influences pharmacokinetic (PK) properties for approximately 40 % of orally administerad drugs (Emami Riedmaier et al. 2020). In cases where the systemic drug exposure is significantly increased (increasing the action of a drug and/or causing unexpected side effects) or decreased (making the drug less effective) in the presence of food a dose adjustment or avoidance of concommittant intake of food and drug is recommended (FDA 2021). According to set criteria, bioequivalence (BE) is reached if ratios between systemic drug exposures in the presence and absence of food are 0.8-1.25.

Authorities recommend that drug developers conduct fed BE studies for orally administered drugs (dosing the drug to individuals in fed and fasted states and evaluating systemic exposures) (FDA 2021). The design of fed BE studies is usually based on *in vitro* dissolution and P_e_ data. Available absorption data and dose size from early clinical studies are also valuable when designing such *in vivo* studies.

The Food Effect PBPK IQ Working Group, consisting of members from 14 major pharmaceutical companies, evaluated the predictive performance of physiologically based PK (PBPK) modelling on food-drug effects (Emami Riedmaier et al. 2020). In their work, they collected and selected 30 drugs with varying effects and sizes of food-drug effects and used physicochemical data (clogD and pK_a_), solubility in aqueous buffer and fasted and fed state simulated intestinal fluids (FaSSIF and FeSSIF), *in vitro* P_e_ (MDCK and Caco-2 cell lines), human *in vivo* P_e_, clinical PK data, and Simcyp and GastroPlus software for modelling/simulating/predicting systemic exposure (area under the plasma concentration-time curve (AUC) maximum plasma concentration (C_max_)) of drugs in the fasted and fed states. Simulated AUC and C_max_ values were compared to corresponding clinically observed estimates. For each compound the work was initiated with the question *“Are fasted C*_*max*_ *and AUC within 0.8-1.25-fold of observed?”*, followed by a decision tree for model verification and optimization, and finally prediction of food effect post optimization. High (within 0.8- to 1.25-fold) to moderate confidence (within 0.5- to 2-fold) was achieved for most of the compounds (15 and 8, respectively). For 7 compounds, prediction confidence was found to be >2-fold (low). For 3 of the compounds, the highly lipophilic amiodarone, danazol and dabrafenib, no results could be generated. Other limitation with the study was that the results were not pure forward-looking predictions and prodrugs and drugs for which absorption was known to be limited by intestinal transporters were excluded.

The ANDROMEDA software by Prosilico for prediction and optimization of human clinical PK (with a major domain for compounds with molecular weight (MW) between 100 and 700 g/mole) has been sucessfully applied and validated for prediction of absorption/biopharmaceutics of small compounds in man in many studies (Fagerholm et al. 2021, 2022, 2023a,b). ANDROMEDA predicted absorption properties, including fraction absorbed (f_a_), efflux by MDR-1, BCRP and MRP2 and *in vivo* dissolution potential (f_diss_), and oral dose, can be used to estimate the size of food interaction effect (ratio between area under the plasma concentration-time curves in the fed (fat food) and fasted states (AUC_fed_/AUC_fast_)).

The main objective was to use ANDROMEDA and actual dose sizes to predict AUC_fed_/AUC_fast_ for small drugs (including those marketed in 2021) and compare results with corresponding measured clinical estimates. Another objective was to compare the performance of ANDROMEDA *vs* PBPK modelling and simulation by The Food Effect PBPK IQ Working Group, where results were generated using Simcyp and GastroPlus software and based on various physicochemical, *in vitro* and *in vivo* data and a decision tree for model verification and optimization.

## Methodology

### Data selection

AUC_fed_/AUC_fast_-data for 29 drugs in Emami Riedmaier et al. (2020) (metoprolol was excluded as food is proposed to influence systemic exposure via changes in blood flows), 17 small drugs marketed in 2021 (maximum 1000 mg oral dose) and 17 other small drugs were selected for this study (Table 1). Clinical food-drug interaction data for drugs not included in the study by Emami Riedmaier et al. were taken from FDA-label documents. Only drugs with MW 100 to 700 g/mole and oral dose of maximally 1250 mg were selected. The selected drugs cover wide ranges of lipophilicity (clogD -2.1 to 7.3), aqueous buffer solubility (0.5 to 128,000 mg/L), human P_e_ (0.1 to 12.5 • 10^−4^ cm/s), f_a_ (very low to complete) and AUC_fed_/AUC_fast_ (0.17 to 5.5).

**Table 1.** The 63 small drugs and their observed AUC_fed_/AUC_fast_, prediction errors with ANDROMEDA and simulation/modelling/prediction error with two PBPK-models (Simcyp and GastroPlus).

| Drug | Observed<br>AUC<sub>fed</sub>/AUC<sub>fast</sub> | ANDROMEDA<br>Prediction<br>error | PBPK Model 1<br>Simulation<br>error | PBPK Model 2<br>Simulation<br>error | Comment |
| --- | --- | --- | --- | --- | --- |
| Alectinib* | 2,92 | <2-fold | 2,54 | <2-fold |  |
| Amiodarone* | 2,36 | <2-fold | >2 (failed) | >2 (failed) | cLogD 6.5 |
| Aprepitant* | 2,81 | 2,66 | not applicable | <2-fold |  |
| Cimetidine* | No food effect | <2-fold | <2-fold | <2-fold |  |
| Clarithromycin* | No food effect | <2-fold | <2-fold | <2-fold |  |
| Dabrafenib* | 0,70 | 2,38 | >2 (failed) | >2 (failed) | cLogD 3.5 |
| Danazol* | 2,90 | <2-fold | >2 (failed) | >2 (failed) | cLogD 7.3 |
| Danirixin* | 0,65 | <2-fold | <2-fold | not applicable |  |
| Etoricoxib* | No food effect | <2-fold | <2-fold | <2-fold |  |
| Fluoxetine* | 0,70 | <2-fold | <2-fold | <2-fold |  |
| Furosemide* | No food effect | <2-fold | <2-fold | <2-fold |  |
| Imatinib* | No food effect | <2-fold | <2-fold | <2-fold |  |
| Isoniazid* | No food effect | <2-fold | <2-fold | <2-fold |  |
| Itraconazole* | 1,66 | <2-fold | <2-fold | <2-fold |  |
| Ivacaftor* | 3,07 | <2-fold | <2-fold | not applicable |  |
| Nefazodone* | No food effect | <2-fold | <2-fold | <2-fold |  |
| Nelfinavir mesylate* | 5,06 | 3,48 | <2-fold | not applicable | 1250 mg dose |
| Nifedipine* | No food effect | <2-fold | <2-fold | <2-fold |  |
| Oseltamivir* | No food effect | <2-fold | <2-fold | not applicable |  |
| Panobinostat* | No food effect | <2-fold | <2-fold | <2-fold |  |
| Pazopanib* | 2,45 | <2-fold | 7,66 | 2,53 |  |
| Phenytoin* | No food effect | <2-fold | <2-fold | <2-fold |  |
| Sotalol* | No food effect | <2-fold | <2-fold | not applicable |  |
| Telaprevir* | 3,71 | 2,66 | 2,36 | <2-fold | 750 mg dose |
| Tezacaftor* | No food effect | <2-fold | <2-fold | not applicable |  |
| Trospium* | 0,17 | 4,13 | <2-fold | 3,40 | Very low P<sub>e</sub> |
| Venetoclax* | 3,41 | <2-fold | not applicable | <2-fold |  |
| Zidovudine* | No food effect | <2-fold | <2-fold | not applicable |  |
| Ziprasidone* | 2,01 | <2-fold | not applicable | <2-fold |  |
| Albendazole | 0,38 | 3,94 | not applicable | not applicable | Low f<sub>diss</sub> |
| Asciminib | 5,02 | 2,16 | not applicable | not applicable |  |
| Atavaquone | 3,30 | <2-fold | not applicable | not applicable |  |
| Atogepant | No food effect | <2-fold | not applicable | not applicable |  |
| Avacopan | 1,72 | <2-fold | not applicable | not applicable |  |
| Belumosudil | 2,00 | <2-fold | not applicable | not applicable |  |
| Belzutifan | No food effect | <2-fold | not applicable | not applicable |  |
| Bicalutamide | No food effect | <2-fold | not applicable | not applicable |  |
| Carvedilol | No food effect | <2-fold | not applicable | not applicable |  |
| Clopidogrel | No food effect | <2-fold | not applicable | not applicable |  |
| Didanosine | 0,53 | <2-fold | not applicable | not applicable |  |
| Etretinate | 5,50 | 2,21 | not applicable | not applicable |  |
| Fenretinide | 3,23 | <2-fold | not applicable | not applicable |  |
| Finasterine | No food effect | <2-fold | not applicable | not applicable |  |
| Finerenone | No food effect | <2-fold | not applicable | not applicable |  |
| Imipramine | No food effect | <2-fold | not applicable | not applicable |  |
| Infigratinib | 2,00 | <2-fold | not applicable | not applicable |  |
| Letrozole | No food effect | <2-fold | not applicable | not applicable |  |
| Maribavir | No food effect | <2-fold | not applicable | not applicable |  |
| Melphalan | 0,45 | <2-fold | not applicable | not applicable |  |
| Mobocertinib | No food effect | <2-fold | not applicable | not applicable |  |
| Penicillamine | 0,49 | <2-fold | not applicable | not applicable |  |
| Ponesimod | No food effect | <2-fold | not applicable | not applicable |  |
| Praziquantel | 3,98 | 2,55 | not applicable | not applicable |  |
| Samidorphan | No food effect | <2-fold | not applicable | not applicable |  |
| Sotorasib | No food effect | <2-fold | not applicable | not applicable |  |
| Tepotinib | 1,60 | <2-fold | not applicable | not applicable |  |
| Terfenadine | No food effect | <2-fold | not applicable | not applicable |  |
| Tivozanib | No food effect | <2-fold | not applicable | not applicable |  |
| Umbralisib | 1,61 | <2-fold | not applicable | not applicable |  |
| Valproic acid | No food effect | <2-fold | not applicable | not applicable |  |
| Vardenafil | No food effect | <2-fold | not applicable | not applicable |  |
| Vericiguat | 1,44 | <2-fold | not applicable | not applicable |  |
| Viloxazine | No food effect | <2-fold | not applicable | not applicable |  |

\*Taken from Emami Riedmaier et al. (2020) and used for method comparison.

### In silico predictions using ANDROMEDA and dose sizes

ANDROMEDA by Prosilico was used to predict absorption parameters (including, for example, f_diss_ and efflux-substrate specificities) and then (based on these data, algorithms and oral dose sizes), AUC_fed_/AUC_fast_. In order to produce true forward-looking predictions, selected drugs were not included in training sets of underlying prediction models. Predicted AUC_fed_/AUC_fast_ with ANDROMEDA has a maximum of 6-fold.

### PBPK modelling, simulation and prediction using Simcyp and GastroPlus

Results were taken from the work by The Food Effect PBPK IQ Working Group (Emami Riedmaier et al. 2020). The group collected and selected drugs (excluding prodrugs and drugs for which absorption was known to be limited by intestinal transporters were excluded) with varying effects and sizes of food-drug effects and used physicochemical data (clogD and pK_a_), solubility in aqueous buffer, FaSSIF and FeSSIF), *in vitro* P_e_ (MDCK and Caco-2 cell lines), human *in vivo* P_e_, clinical PK data, and Simcyp (version 7.1) and GastroPlus (version 9.5) software for modelling/simulating AUC_fast_, AUC_fed_ and AUC_fed_/AUC_fast_. For each compound, the work was initiated with the question *“Are fasted C*_*max*_ *and AUC within 0.8-1.25-fold of observed”*, followed by a decision tree for model verification and optimization, and finally, modelling/simulation of food effect post optimization. Results are mainly presented as the combination of models 1 and 2.

## Results & Discussion

### In silico predictions using ANDROMEDA and dose sizes

Results obtained with ANDROMEDA are shown in Figure 1 and Table 1. Predicted AUC_fed_/AUC_fast_ had mean, median and maximum errors of 1.5-, 1.2- and 4.1-fold, respectively, and the predictive accuracy (correlation between predicted and observed AUC_fed_/AUC_fast_; Q^2^) was 0.3. 14 % of predictions had >2-fold error. For 32 % of drugs with AUC_fed_/AUC_fast_ between 0.8 and 1.25, no clinical food interaction was predicted, and for 72 % of drugs, food interaction class (increase, unchanged or decreased AUC) was correctly predicted (Figure 2). 2 % of drugs with AUC_fed_/AUC_fast_>1.25 were incorrectly predicted to have an AUC_fed_/AUC_fast_<0.8, and 2 % of drugs with AUC_fed_/AUC_fast_<0.8 were incorrectly predicted to have an AUC_fed_/AUC_fast_>1.25. Every predicted value >2.2 corresponded to a clinically significant food-drug interaction (AUC_fed_/AUC_fast_≥1.25).

**Figure 1.**
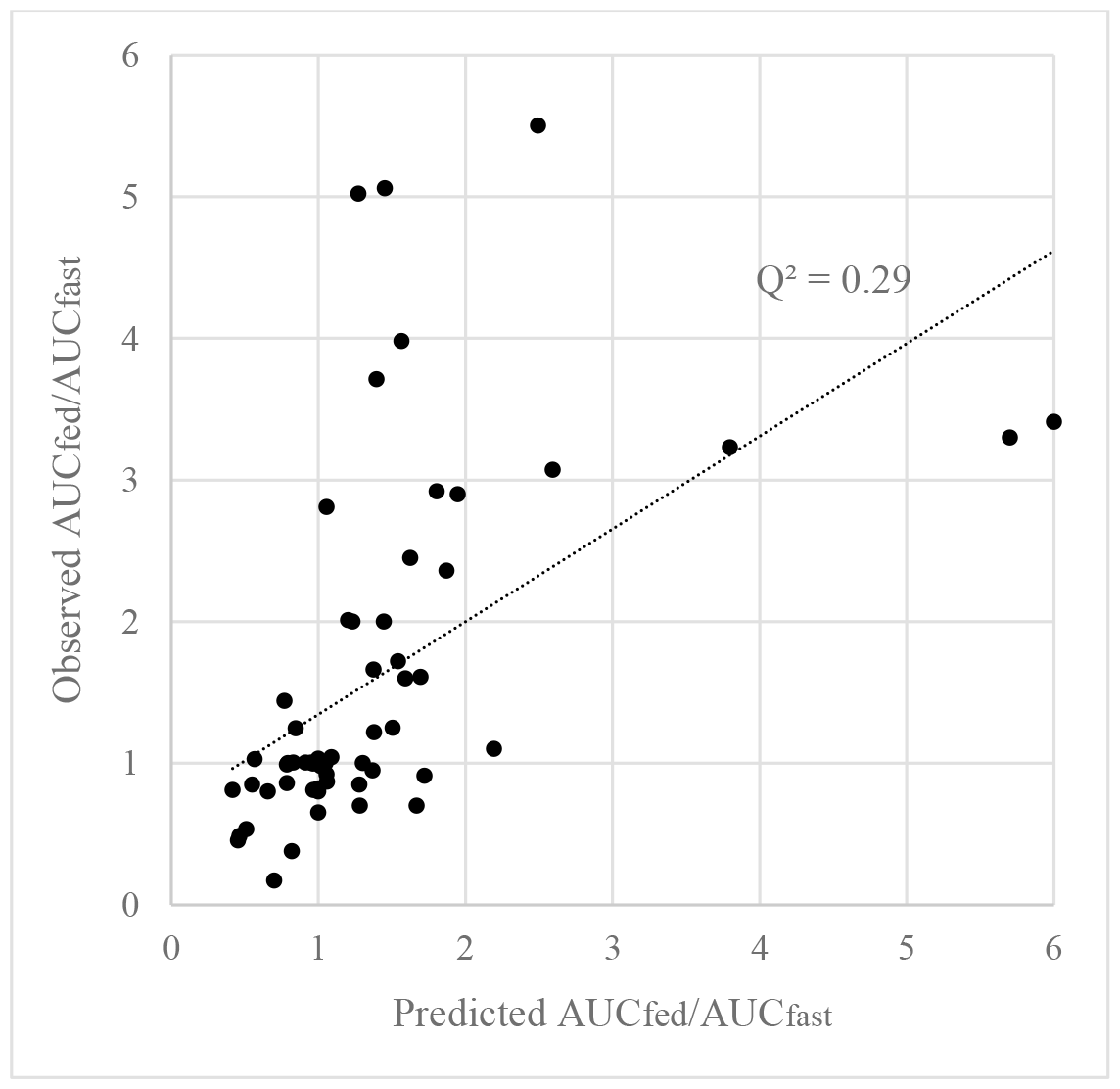
The correlation between predicted (ANDROMEDA) and observed AUC_fed_/AUC_fast_ for the 63 small drugs.

**Figure 2.**
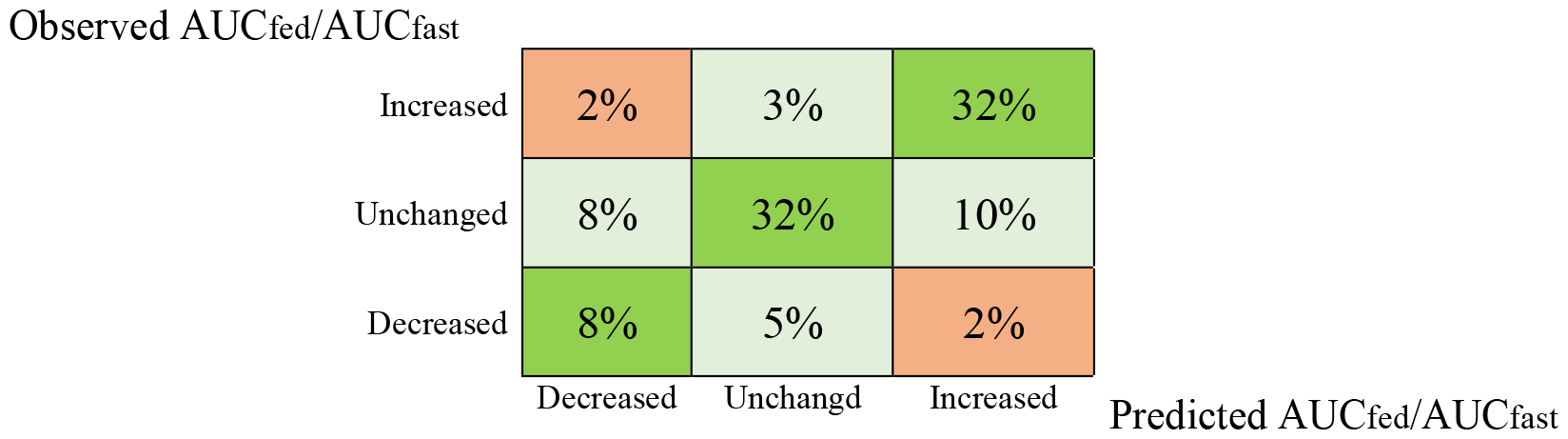
Predicted *vs* observed AUC_fed_/AUC_fast_-classes (%) for the 63 drugs. Decreased when AUC_fed_/AUC_fast_<0.8; unchanged when AUC_fed_/AUC_fast_=0.8-1.25; increased when AUC_fed_/AUC_fast_>1.25.

The largest prediction errors were found for albendazole (3.9-fold underprediction; enhanced AUC_fed_/AUC_fast_ observed and predicted; low f_diss_ and f_a_), nelfinavir mesylate (3.5-fold underprediction; enhanced AUC_fed_/AUC_fast_ observed and predicted; 1250 mg dose*) and trospium (4.1-fold overprediction; reduced AUC_fed_/AUC_fast_ observed and predicted; low P_e_ and f_a_). Despite prediction errors of this size, none of these 3 compounds showed incorrectly predicted AUC_fed_/AUC_fast_-class.

*There is a trend towards slightly higher prediction errors at higher oral doses.

The predictive performance was enhanced with introducing actual dose size. 5 to 40 % higher mean, median and maximum prediction errors were reached when not considering dose size.

### PBPK modelling and simulation using Simcyp and GastroPlus

Predicted AUC_fed_/AUC_fast_ had mean, median and maximum errors of 1.5-, 1.2- and 7.7-fold, respectively (n=26; excluding the non-quantifiable amiodarone, danazol and dabrafenib), and the predictive accuracy (correlation between simulated and observed AUC_fed_/AUC_fast_; R^2^) was 0.4 (Figure 3; Table 1). 12 % of predictions had >2-fold error (23 % when including amiodarone, danazol and dabrafenib). For 77 % of drugs with AUC_fed_/AUC_fast_ between 0.8 and 1.25, no clinical food interaction was predicted, and for 81 % of drugs (73 % when including amiodarone, danazol and dabrafenib), food interaction class (increase, unchanged or decreased AUC) was correctly predicted.

**Figure 3.**
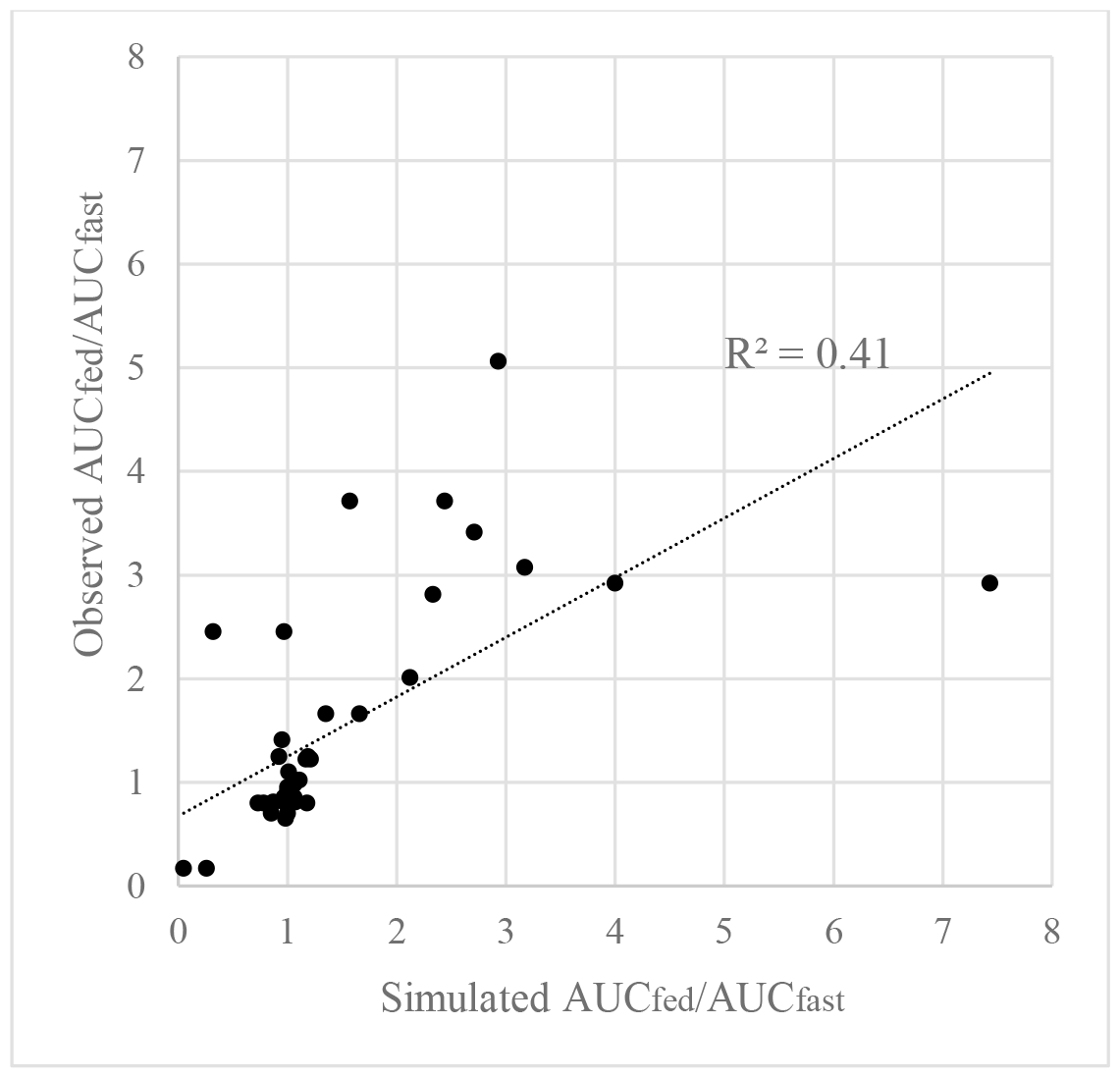
The correlation between simulated (PBPK; Simcyp and GastroPlus) and observed AUC_fed_/AUC_fast_ for 26 small drugs. Amiodarone, danazol and dabrafenib with non-quantifiable results were not included.

One drug (3 %) with AUC_fed_/AUC_fast_>1.25 was incorrectly simulated to have an AUC_fed_/AUC_fast_<0.8.

PBPK model 2 had better performance than PBPK model 1, with higher R^2^ (0.7 *vs* 0.3), lower mean prediction error (1.4- vs 1.6-fold) and lower maximum prediction error (3.4- *vs* 7.7-fold). When assuming a prediction error of 4-fold for amiodarone, danazol and dabrafenib R^2^-values decreased to ca 0.2-0.3.

No *in vitro* P_e_-data was available for 11 (38 %) of the 29 compounds, including all with cLogD≥4.8. This is a limitation for *in vitro* to *in vivo*-prediction of food effects, especially for highly lipophilic drugs.

A R^2^ of 0.2 was found for log FeSSIF/FaSSIF-solubility *vs* AUC_fed_/AUC_fast_, indicating a weak impact of fed *vs* fast solubility using simulated intestinal fluids.

### Comparing ANDROMEDA and PBPK (Simcyp and GastroPlus)

Overall, ANDROMEDA and PBPK showed similar results. Mean, median and maximum errors for PBPK and ANDROMEDA were 1.5-, 1.2- and 7.7-fold (higher if including 3 non-predictable compounds) and 1.5-, 1.2- and 4.1-fold, respectively, the R^2^ was 0.4 (ca 0.2-0.3 if including 3 non-predictable compounds) and the Q^2^ was 0.3, 12 % (23 % if including 3 non-predictable compounds) and 14 % of predictions with >2-fold error, respectively, and 81 % (73 % if including 3 non-predictable compounds) and 72 % of drugs with correctly predicted food interaction class.

For the same set of 26 compounds as in the PBPK studies, ANDROMEDA reached 1.7-, 1.5- and 4.1-fold mean, median and maximum prediction errors, respectively, a Q^2^ of 0.23 (same as with amiodarone, danazol and dabrafenib included) and 59 % correct classifications.

The PBPK models seem to simulate/model/predict AUC_fed_/AUC_fast_ comparably good for compounds with no or minor food interaction effects. This might be due to inclusion of such knowledge into the decision tree (*“Are fasted C*_*max*_ *and AUC within 0.8-1.25-fold of observed?”*) for modelling and simulations. Exclusion of compounds without food effect increased mean errors (1.5- to 1.7-fold for both ANDROMEDA and PBPK) and reduced Q^2^ (to 0.24 for ANDROMEDA predictions) and R^2^ (0.28 for PBPK simulations).

Major advantages with ANROMEDA include that results are produced directly from molecular structure and oral dose, without *in vitro* and *in vivo* PK and AUC_fed_/AUC_fast_ data. The level of predictive accuracy was overall similar to results obtained with laboratory and clinical data, PBPK and simulations and modelling, however, with lower maximum error and higher compound coverage. In addition, ANDROMEDA was evaluated based on a dataset with greater maximum food effect (than for PBPK) and with compounds with significant contribution by active intestinal transport (these were said to be excluded in the PBPK-study).

## Conclusion

The results validate ANDROMEDA for prediction of fat food-drug interaction size for small drugs in man. Major advantages with the methodology include that prediction results are produced directly from molecular structure and oral dose and are similar to PBPK-simulation results obtained using *in vitro* and clinical data. Furthermore, ANDROMEDA showed lower maximum errors and wider compound range (covering compounds with the highest lipophilicity).

